# Disorder mediated oligomerization of DISC1 proteins revealed by coarse-grained computer simulations

**DOI:** 10.1101/660910

**Authors:** Julien Roche, Davit A Potoyan

**Affiliations:** Department of Chemistry, Iowa State University Ames IA; Department of Biochemistry and Molecular Biology, Iowa State University Ames IA

**Keywords:** Chromatin, Eukaryotic Nucleus, Meso-scale modeling, phase separation

## Abstract

Disrupted-in-Schizophrenia-1 (DISC1) is a scaffold protein of significant importance for neurodevelopment and a prominent candidate protein in the etiology of mental disorders. In this work, we investigate the role of conformational heterogeneity and local structural disorder in the oligomerization pathway of the full-length DISC1 and of two truncation variants. Through extensive coarse-grained molecular dynamics simulations with a predictive energy landscape based model, we reveal the general mechanistic principles of DISC1 oligomerization. We found that both conformational heterogeneity and structural disorder play an important role in the dimerization pathway of DISC1. This study sheds light on the differences in oligomerization pathways of the full-length protein compared to the truncated variants produced by a chromosomal translocation associated with schizophrenia.

## Introduction

Regulation, recognition and cell signaling involve the co-ordinated actions of many players, including scaffold proteins that provide selective spatial and temporal coordination among interacting proteins. Scaffold proteins have typically no catalytic activity but instead bring together interacting proteins within a signaling pathway to facilitate or modify the specificity of their interactions. A common characteristic of scaffold proteins is the extent of structural disorder(1). Structural disorder is intrinsically associated with several key attributes of scaffold proteins, such as the ability to bind multiple partners, mediating transient interactions and undergoing a complex array of regulatory post-translational modifications(2).

To understand the role of structural disorder in mediating the function of scaffold proteins, we carried out extensive molecular simulations on DISC1, a canonical scaffold protein involved in neuron differentiation and migration. *DISC1* was originally identified at the break-point of a (1:11) chromosomal translocation in a Scottish family with a high loading of major mental illness (3, 4) and is now established by considerable genetic evidence as a risk factor for a wide range of psychiatric disorders (5–8). Beyond the balanced chromosomal translocation, multiple other sequence variations of *DISC1* are associated with schizophrenia, depression and autism (9, 10) The product of *DISC1* is a scaffold protein that regulates the activity, mostly through inhibition, of a considerable number of enzymes of interrelated signaling cascades in the central nervous system (11–13). Despite growing appreciation of its role in the etiology of mental disorders, almost nothing is known about the structural properties of DISC1 by itself and in complex with its protein partners. DISC1 is a 854-residue long protein that shares many features with other long scaffold proteins containing fully or partially disordered regions. Bioinformatics predictions suggest that the first 350 amino acids of DISC1 are predominantly disordered (Fig S1) whereas the remainder of the protein is rich in a-helical and coiled coil motifs. The central region, which is essential for DISC1 self-association, is predicted to form a discontinuous coiled-coil domain with no continuous stretches greater than 50 amino acids (14). An early study has predicted the presence of two UVR domains in this region, consisting of two a-helices, packed against each other in an anti-parallel hairpin fold (15). The C-terminus contains two regions with coiled-coil propensity and two leucine zippers motifs (14). Zhang and coworkers have recently solved the atomic structure of DISC1 C-terminal tail (aa. 765-835) in complex with Nde1, revealing a helix-turn-helix structure (16), which was later confirmed by SAXS (17). Overall, the C-terminal region appears to possess the characteristics of a series of helical bundles that mediate protein-protein interactions.

Wagner and coworkers have shown that the full-length protein exists in equilibrium between octamers and dimers but this equilibrium is delicate and examples have been reported where single point mutations or truncations affect the oligomerization state of DISC1 (18–20). The oligomerization state of DISC1 appears to be a delicate equilibrium between dimeric and octameric species, potentially modulated by PKA-induced phosphorylation (21). Disease associated mutations such as S704C have also been reported to affect the oligomerization state of DISC1 (18). Yet, the structure of DISC1 oligomers and pathways to oligomerization remain largely unknown.

To unravel the specific contributions of conformational dynamics and structural disorder to the formation of oligomers, we conducted extensive long-time scale simulations using coarse-grained predictive model of the full-length DISC1 and the truncated fragments resulting from the (1:11) chromosomal translocation associated with schizophrenia (3, 4). Through a set of constant temperature simulations we first explored equilibrium conformations ensemble of all monomeric units revealing large differences in the conformational ensembles sampled by the full-length protein and the N- and C-fragments. We then ran a distinct set of constant temperature simulations and umbrella sampling simulations to study a set of candidate oligomerization structures and their oligomerization free energy landscapes. By employing various measures for protein contact analysis in the form of contact frequency maps, intra-domain contain distributions, and principal component analysis in the space of residue-residue contacts we have revealed interplay of contacts that shapes distinct conformations preferences and binding modes of subunits.

## Methods

We used Phyre2 (22) to create an initial 3D model of the full-length DISC1 based on homology modeling. This initial structural model (691 amino acids over a total of 854 residues were modelled at >90% confidence) was further refined with ModRefiner (23) and used for extended MD simulations with the coarse-grained protein force field AWSEM (Associative memory, Water mediated, Structure and Energy Model) (24). The Hamiltonian of AWSEM combines both knowledge based terms acting at local secondary structure level and physics based terms such as hydrogen bonding and water-mediated interactions for capturing interactions at the level of tertiary structures(25). The judicious combination of physical and knowledge based terms has been shown to perform well for *de novo* predictions of majority of proteins with foldable landscapes including native states and allosteric conformational states of single and multi-domain proteins (25–27). Thanks to the predictive nature of AWSEM model it has found wide applicability for solving various biophysical problems including predictions of monomeric protein units, protein aggregation (28, 29), protein-DNA assembly (30, 31), proteins with unstructured regions (32) and recently also for natively disordered proteins (33). In this work we have used the AWSEM Hamiltonian supplemented with electrostatic potential (34) and multiple fragment memories(32, 35) to account for disordered nature of the DISC1 sequence.

All of the molecular dynamics simulations were ran using the LAMMPS molecular dynamics simulator. The Langevin integrator in LAMMPS was employed with a 5*fs* integration time step and with a damping time of 10^3^*fs*. Simulations were run for 20 − 50 million steps until convergence was reached which was checked by comparing different sets of independent runs for agreements between first few eigenvectors of contact-based principal components. Conformations of monomers and dimers were saved at 5000 intervals. For *de novo* prediction of dimers we have initiated monomers in parallel and anti-parallel orientation and carried out short simulated annealing runs followed by constant temperature simulations at *T* = 300*K*. Umbrella sampling simulations were initiated using conformations generated from monomer and C-,N-terminal fragment simulations. We have considered both parallel and anti-parallel orientations. Center of mass harmonic constraint *U*_*i*_ = *k*_*com*_(*R*_*com*_ − ⟨*R*_*i*_ ⟩)^2^ have been applied to the groups of residues spanning the central regions of the DISC1 (Fig. 1) in the case of full length monomer and all residues in the case of N and C-terminal fragments. We used *k*_*com*_ ∼ 0.12*kcal/*Å^2^ for spring constants and 50, Δ⟨*R*_*i*_⟩ = 2Å separated windows which were generated from the bound states. Weighted Histogram Analysis Method (WHAM) has been used for reconstructed unbiased estimates of free energy as a function of center of mass coordinate *F* (*R*_*com*_).

**Fig. 1.**
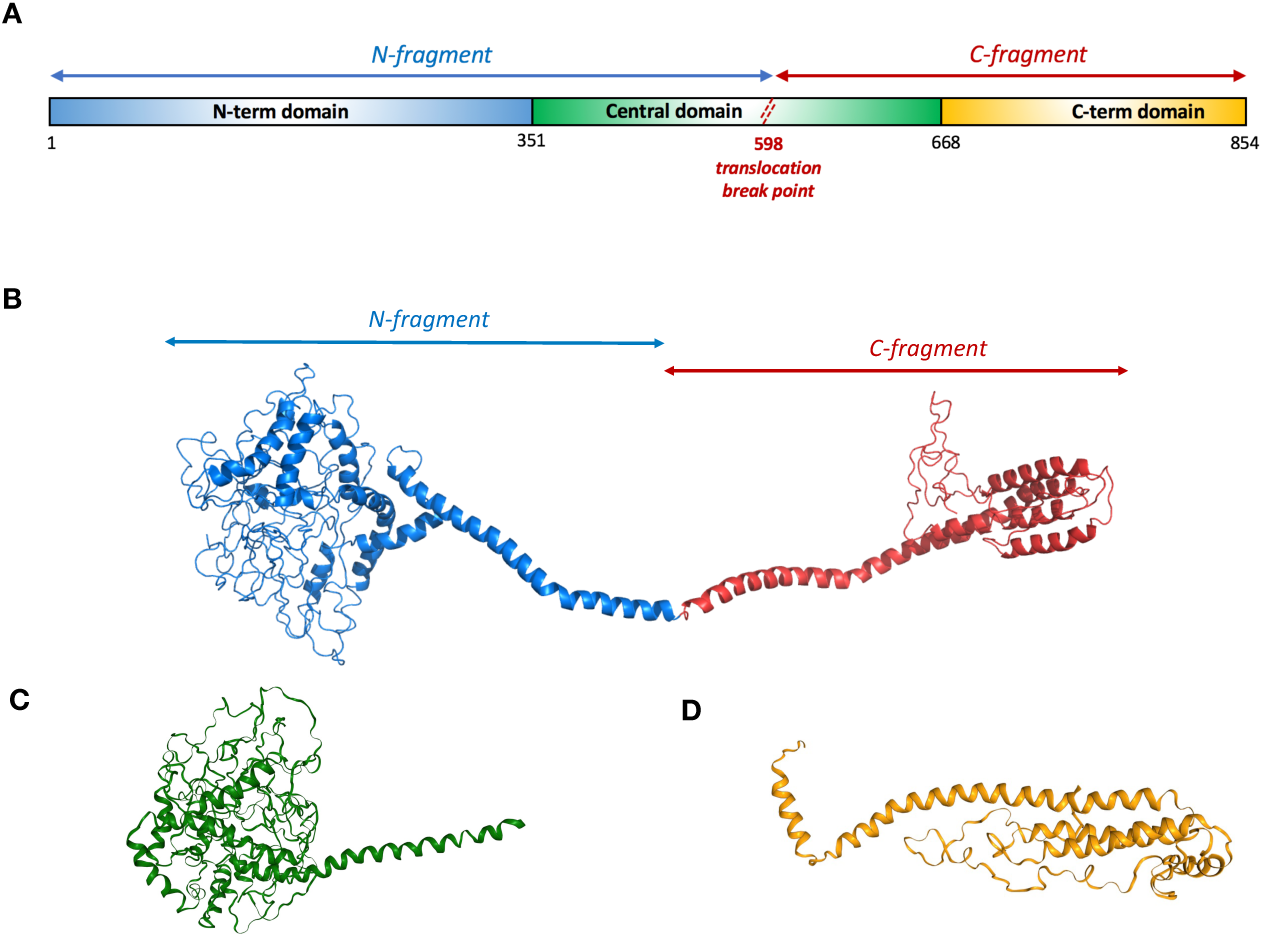
Shown are the domain organization of (A) full length, (B) N-terminal and (C) C-terminal fragments of DISC1 sequence together with representative structures obtained through molecular dynamics simulations. Consistent color coding of structural domains is used throughout the manuscript with blue/green corresponding to N-terminal domain(s) and red/orange to C-terminal domain(s).

For the contact analysis we have considered *C*_*α*_ atoms excluding contacts between (*i, j*) pairs of residues that are |*i* − *j*| *>* 3 distance away in sequence space. We have used distances satisfying *d*_*ij*_ *<* 8Å condition as the definition of contacts. With this definition of contacts we have carried out principal component analysis that reveals the non-local contact making and breaking in conformational ensembles of monomers and oligomers within first few principal components.

## Results and Discussion

### A. In its monomeric form DISC1 is in equilibrium between extended and compact states

To fully describe the mechanisms of oligomerization of DISC1, we first examined the conformational ensemble populated by the monomeric full-length protein. We achieved an extensive sampling of conformations through 20 independent simulations that were initiated by performing simulated annealing runs followed by more than 200*µs* simulations at constant temperature *T* = 300*K*. The conformational ensemble generated shows an heterogeneous distribution of mostly helical structures, with significant structural disorder localized within N-terminal domain (Fig 1,2), in good agreement with with bioinformatic secondary structure predictions (Fig, S1). To further characterize the degree of heterogeneity and structural disorder in DISC1 ensemble, we employed two separate measures; radius of gyration and contact-metrics (contact frequency map, contact based PCA) for global and local scales characterization respectively. In the absence of well defined 3D structures, the radius of gyration has been a central quantity for characterizing the extent of structural disorder in proteins. However, in situations where there is significant secondary structural content present, global measures of disorder such as radius of gyration, end-to end distance, structure factor are less informative. Contact based measures have the advantage of revealing more detailed picture of disorder throughout sequence as well as revealing dynamic and thermodynamic driving forces that shape energy landscapes of disordered proteins.

**Fig. 2.**
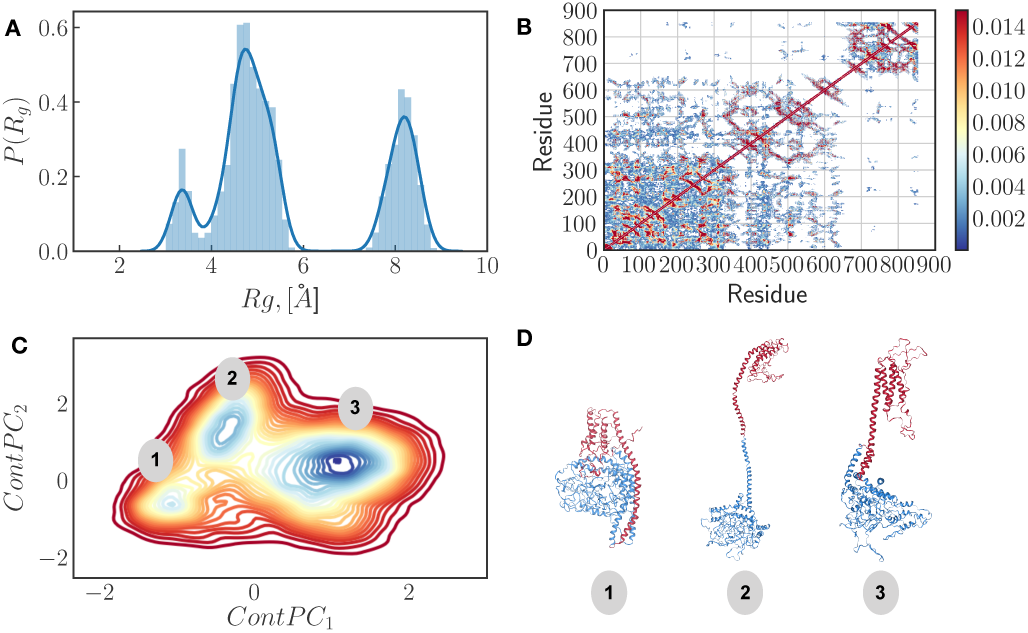
Shown are (A) Distribution of radii of gyration for full monomer unit of DISC1 (B) Contact frequency map showing the *C*_*α*_ − *C*_*α*_ contacts that are observed more frequently in simulations (C) Probability distribution of simulated trajectories onto principal components in the space of physical *C*_*α*_ − *C*_*α*_ contacts that correspond to high contact frequencies (D) Representative structures from simulations corresponding to distinct regions of the first two principal components.

The distribution of radius of gyration clearly shows the presence of at least three major populations, with average radius of gyrations of 3.5, 4.6 and 8.2 A (Fig. 2A). Analyzing frequency of contact formation in DISC1 ensemble we found specific pattern of contacts within the N- and C-terminal domains while the central domain shows largely smeared contacts with the N-terminal domain (Fig. 2B). We then performed a principal component analysis (PCA) of the distances of statistically significant contact between residue pairs (p > 0.001) identified in the frequency map. The first principal components identify sets of contacts which contribute the most to the variance and these pairs typically correlate with more global structural changes. The probability distribution of conformational ensemble onto the first two contact principal modes shows indeed that the high variance contacts between the central and N-terminal domains are significant drivers of observed conformational heterogeneity (Fig. 2C). In good agreement with the distribution of radius of gyration, we found with the principal component analysis three major populations within the conformational ensemble of DISC1, corresponding to compact, semi-compact and extended states (labeled 1, 2 and 3 respectively in figure 2C and 2D).

### B. The N- and C-terminal fragments are conformationally less heterogeneous than the full-length DISC1 but not less disordered

Examining the N-terminal fragment of DISC1 (residue 1-597, see figure 1C) we found drastically different conformational behaviour. The radius of gyration distribution shows largely uni-modal distribution in this case as opposed to well defined multi-modal features of full length DISC1 (Fig. 3A). The decrease in conformational heterogeneity however does not imply decrease in disorder. The map of contact frequencies shows this very clearly. Indeed, we observed an increase in local disorder which is likely driven by the absence of contacts formed between the N-terminal and central domains (Fig. 3B). The principal component analysis of statistically significant contact making residue pairs (p > 0.001) revealed that contact making and breaking do not drive any large scale coherent structural changes (Fig. 3C). Instead, contact making and breaking is highly delocalized and irregular as one would expect from protein with significant amount of disorder. Overall, the conformational ensemble of the N-terminal fragment is relatively homogeneous, with a single major population corresponding a compact and largely discorded state (Fig. 3D).

**Fig. 3.**
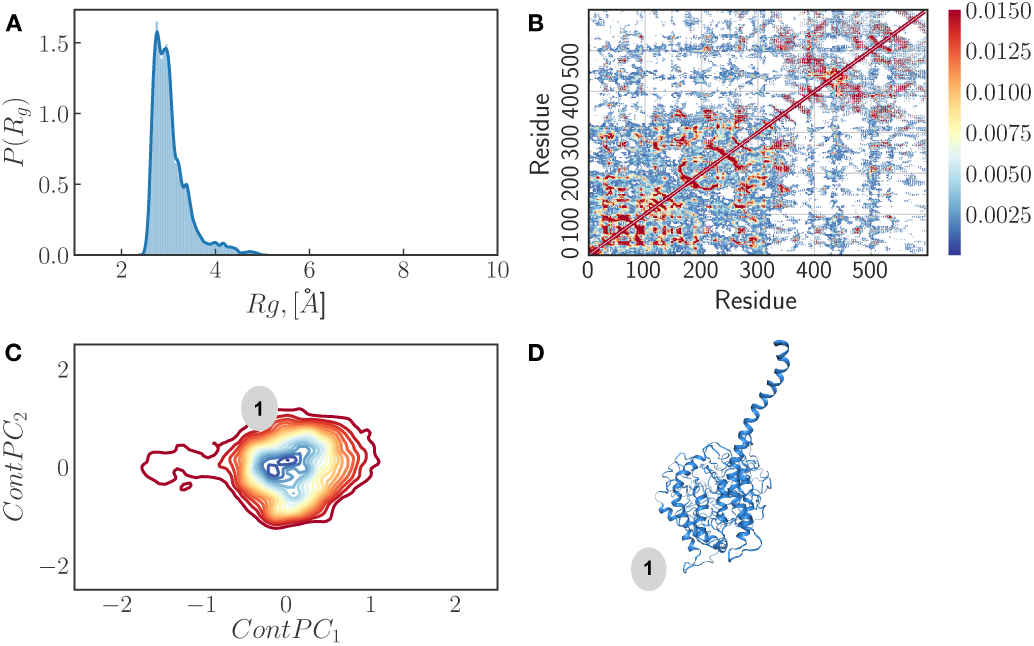
Shown are (A) Distribution of radii of gyration for N-terminal fragment unit of DISC1 (B) Contact frequency map showing the *C*_*α*_ − *C*_*α*_ contacts that are observed more frequently in simulations (C) Probability distribution of simulated trajectories onto principal components in the space of physical *C*_*α*_ − *C*_*α*_ contacts that correspond to high contact frequencies (D) Representative structures from simulations corresponding to distinct regions of principal components.

We then examined the C-terminal fragment (residue 598-854, see Figure 1C). We found conformational features for the C-terminal fragment different from the full length DISC1 and N-terminal fragment. In particular we observed much less structural disorder relative to N-terminal fragment yet the radius of gyration distribution shows a non-negligible degree of conformational heterogeneity present (Fig. 4A). This heterogeneity appears to be is driven by frustration to form bundled alpha helices versus the conformational entropy favored elongated forms. Analysis of contact frequency maps shows that the conformational ensemble is populated with relatively rigid local alpha helical segments that can adopt multiple different global folds due to elongated nature of the chain (Fig. 4B). Principal component analysis of statistically significant contacts (p > 0.001) reveals that indeed the most variant set of contacts drive large scale structural changes through various sets of contact making and breaking (Fig. 4C). Based on this analysis, the C-terminal fragment seems to populate an heterogeneous ensemble of helical bundles, with in its most compact form, up to four short helices (Fig. 4D).

**Fig. 4.**
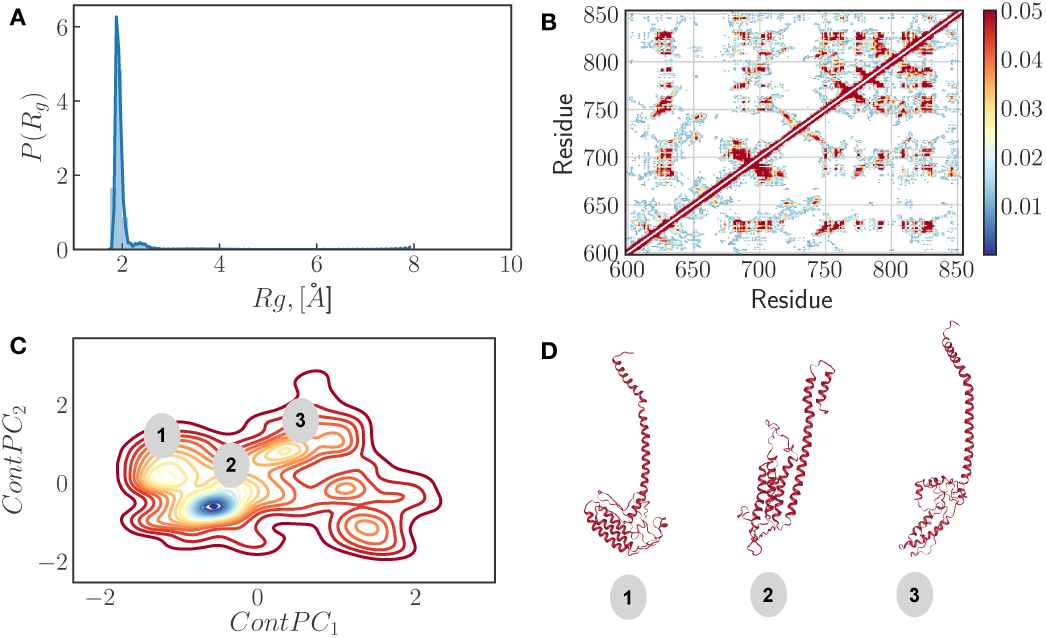
Shown are (A) Distribution of radii of gyration for C-terminal fragment unit of DISC1 (B) Contact frequency map showing the *C*_*α*_ − *C*_*α*_ contacts that are observed more frequently in simulations (C) Probability distribution of simulated trajectories onto principal components in the space of physical *C*_*α*_ − *C*_*α*_ contacts that correspond to high contact frequencies (D) Representative structures from simulations corresponding to distinct regions of principal components.

Overall, the conformational ensemble of the N-terminal and C-terminal fragments are significantly different from that of the full-length DISC1, suggesting that inter-domain contacts participate in the structural heterogeneity of the full-length protein. This interplay between structural heterogeneity on a local and global scale hints at the large conformational space accessible to dimeric structures.

### C. Both conformational heterogeneity and structural disorder mediate the oligomerization of DISC1

Next, we analyze various dimeric combinations of DISC1 and and investigate how conformational heterogeneity and local structural disorder mediate the oligomerization properties of DISC1. Disordered regions in proteins are usually viewed as transient states that can facilitate protein-protein interfaces upon binding (36). The discovery of widespread presence of fuzzy complexes (37–39), however, has shown that the spectrum of action for disordered proteins is much broader (40, 41). Many cases have been uncovered where structural disorder is not only maintained but even generated through protein-protein and protein-nucleic acid interactions (42–44). With DISC1, we found a clear case supporting the role of both conformational heterogeneity and local structural disorder in facilitating the formation of oligomerization interfaces.

To shed light on the complexities of DISC1 oligomerization pathways reported by experiments, we carried out two kinds of simulations aimed at (i) predicting a structural model of the full-length and truncated dimers of DISC1 and (ii) sampling the free energy landscape associated with these oligomerization pathways. For the structural prediction part, we generated two sets of initial 3d models for each construct, representing the parallel and anti-parallel orientation of the sub-units (Fig. 5).

**Fig. 5.**
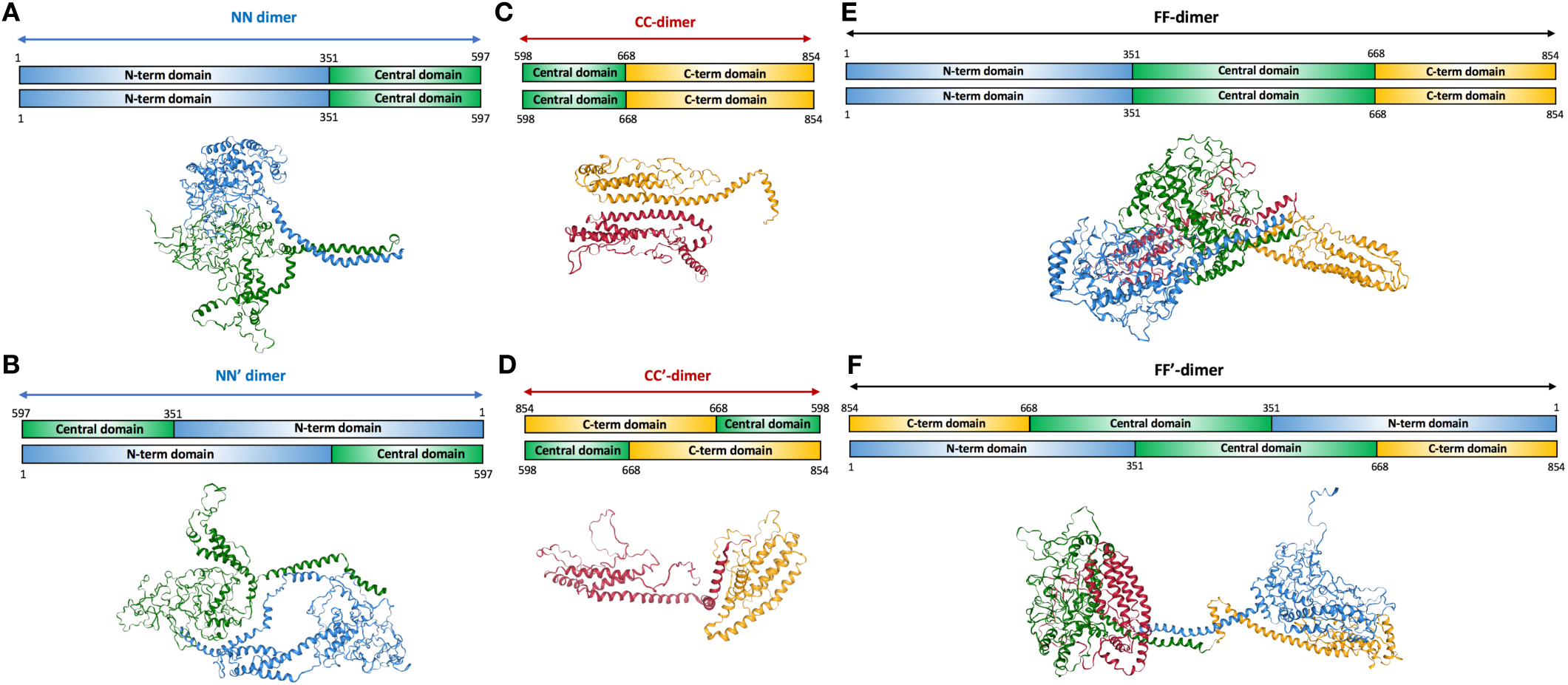
Shown are the various combination of dimeric constructs studied here, with (A) the parallel N-terminal fragment dimer (NN), (B) anti-parallel fragment dimer (NN’), (C) parallel C-terminal fragment dimer (CC), anti-parallel C-terminal fragment (CC’), (E) parallel full-length dimer (FF) and (F) anti-parallel full-length dimer (FF’).

To generate ensemble of dimer states we first ran short time simulated annealing steps followed by long time scale constant temperature molecular dynamics simulations. We found that the dimeric models initiated from extended conformations of the full length DISC1 lead to both extended and collapsed dimers corresponding to parallel and anti-parallel orientations respectively. Furthermore dimerization is accompanied by large scale rearrangement of intra and inter-domain contacts (Fig S2). Dimerization of the N- and C-terminal fragments likewise is followed by large structural rearrangements within each subunit, which are driven by the disruption of intra-domain contacts and formation of inter-domain contacts. We quantified the interplay between intra- and inter-domain contacts by computing the fraction of contacts formed within and between monomeric vs dimeric units. The fraction of contacts is evaluated by computing the ratio *n*_*cont*_(*t*)*/N*_*pairs*_, where *n*_*cont*_(*t*) is the number of *C*_*α*_ − *C*_*α*_ distances which satisfy the *r*_*ij*_ *<* 8Å condition for every state in the sampled conformational ensemble indexed by time co-ordinate *t* (multiple equilibrium trajectories have been combined together). The *N*_*pairs*_ is the total number of *C*_*α*_ − *C*_*α*_ pairs, which provides a normalization measure for comparing fragments vs monomers and dimers (Fig. 6).

**Fig. 6.**
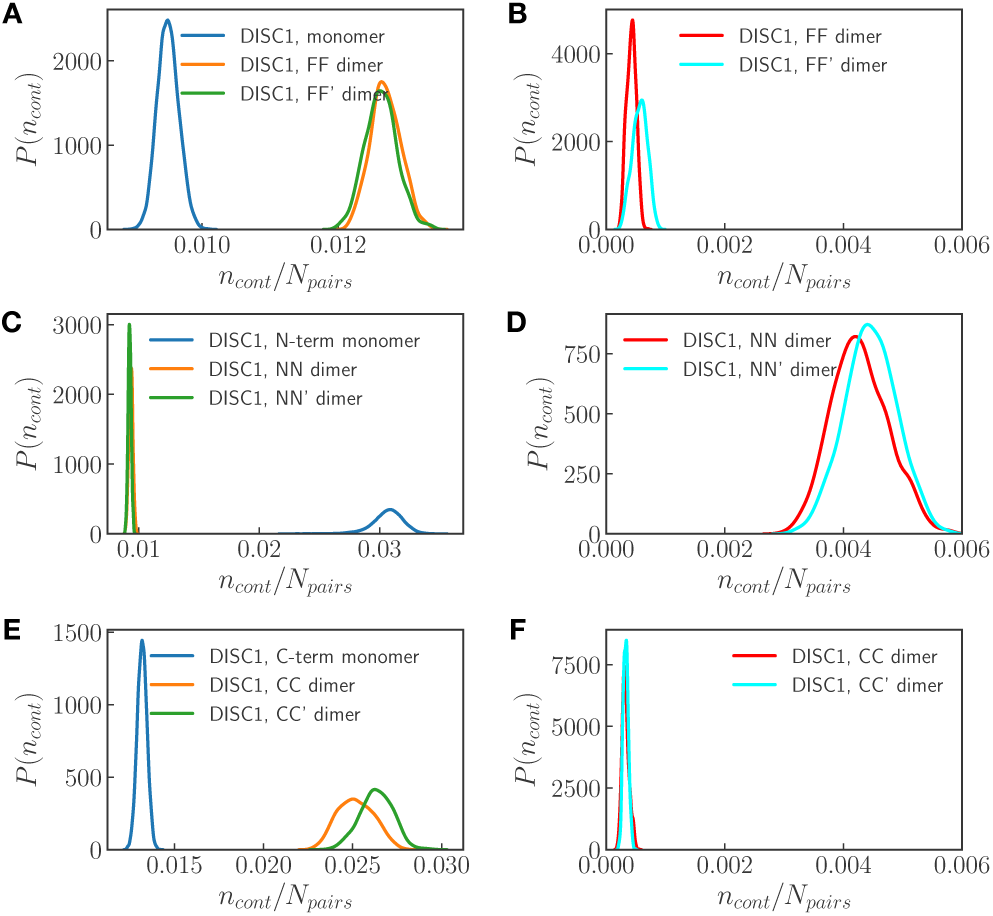
Shown are distribution of *C*_*α*_ − *C*_*α*_ contacts evaluated for (A) intra-chain contacts of full length DISC1 monomer and dimer units. (B) inter-chain contacts for full length DISC1 dimer units. (C) intra-chain contacts of N-terminal DISC1 monomer and dimer units. (D) inter-chain contacts for DISC1 N-terminal fragment dimer units. (E) intra-chain contacts of C-terminal DISC1 monomer and dimer units. (F) inter-chain contacts for DISC1 C-terminal fragment dimer units.

Examining the intra-domain contact distributions for the full length DISC1, we found that dimerization results in an enhancement of contact formation withing each subunit relative to the monomeric form (Fig. 6A). This can be explained by the fact that conformational ensemble of DISC1 monomer includes various collapsed, exposed and intermediate forms which inter-convert via extensive contact making/breaking transitions whereas in the dimeric form, subunits are locked into one major conformational state. We observed no significant difference in terms of intra domain contact distribution between parallel (FF) and anti-parallel dimeric forms (FF’). The distribution of inter-domain contacts (Fig. 6B), however, reveals that dimers with anti-parallel orientation (FF’) form more contacts, due to their collapsed nature, than dimer with parallel orientation (FF’). These results suggest that the flexible nature of the DISC1 manifests not only in conformational heterogeneity at the monomeric level but also in the dimeric form.

In the case of the dimers formed by the N-terminal fragment of DISC1 (NN and NN’), we found that dimerization results in a loss of intra-domain contacts relative to the free monomeric unit(Fig. 6C). This suggests that the formation of dimer interface in the N-terminal region is the result of a trade off between inter- and intra-domain contacts. We observed again no significant different in terms of intra-domain contact distribution between the parallel (NN) and anti-parallel dimeric forms (NN’). The distribution of inter-domain contacts (Fig. 6D), however, reveals that dimers with anti-parallel orientation of N-terminal domain (NN’) are forming a more extensive network of contacts relative to its parallel counterpart (NN). These observations suggest that structural disorder in the N-terminal region is a significant driver of oligomerization by promoting the formation of an extensive network of inter-domain contacts.

For dimers formed by the C-terminal fragment of DISC1 (CC and CC’), we found that the distribution of intra-domain contacts is similar to that of the full length DISC1 (Fig. 6E), suggesting that dimerization of the C-terminal region results in large gain of intra-domain contacts, which is enough to offset the loss of contacts by the N-terminal fragment upon formation of full-length dimers. The distinction in terms of intradomain contact distribution between parallel (CC) and anti-parallel dimeric forms (CC’) appears again negligible. The extent of inter-domain contacts (Fig. 6F) is notably smaller that of the N-terminal fragment and full length DISC1 (keeping in mind that *N*_*pairs*_ ∼ *L*^2^ with *L* being sequence length) which can be rationalized by the absence of structural disorder in the C-terminal region.

Finally, to characterize the oligomerization propensity of the various dimeric conformational states described above, we have computed the free energy profiles of dimerization (Fig. 7) for each putative dimer by carrying out umbrella sampling simulations with an harmonic bias applied to both sub-units. The free energy profiles obtained for the full dimer initiated from the extended monomeric state (structure 2 in Fig. 2D) show that the parallel (FF) and anti-parallel conformations (FF’) have drastically different propensities for dimerization (Fig. 7A). For reference we have also calculated the free energy profiles of association dimers initiated from the collapsed monomeric state (structure 3 in the Fig. 2D). We observed that such dimers maintain their collapsed states, suggesting that in the absence of exposed dimerization surfaces, the free energy gain is significantly diminished (Fig. 7B). Such large free energy gap between collapsed and extended dimeric forms also suggests that both conformational heterogeneity and structural disorder, which modulated the extent of exposed surface area, is important for the formation of stable dimers.

**Fig. 7.**
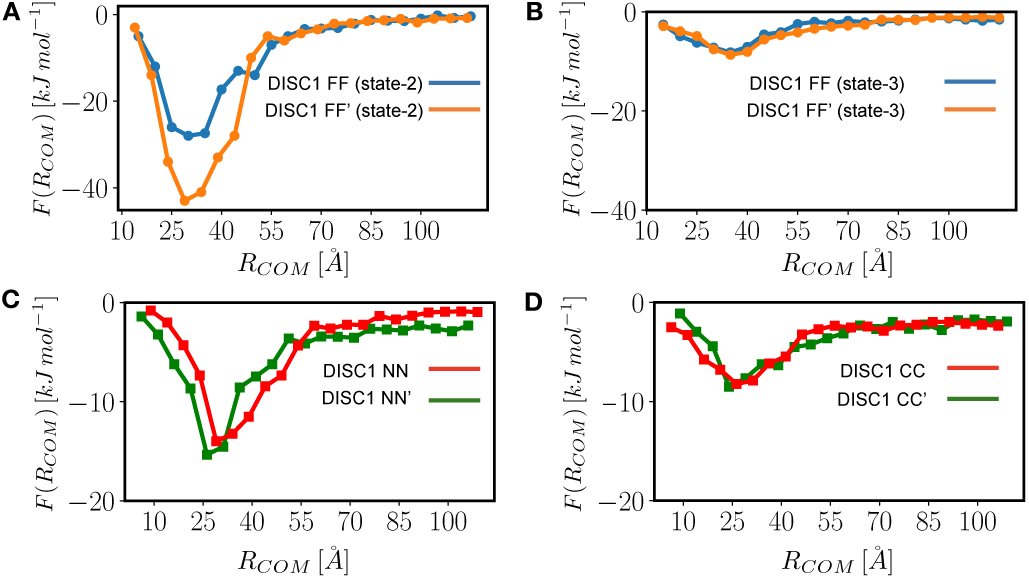
Shown are free energy profiles of binding/unbinding of the different DISC1 dimers with parallel/anti-parallel orientations. (A) Free energy of dimerization with parallel and anti-parallel orientations of full length sequence (B) Free energy profile of binding between collapsed states of the full length sequence. (C) Free energy profile for N-terminal fragment dimerization with parallel and anti-parallel orientation (D) Free energy profile for C-terminal fragment dimerization with parallel and anti-parallel orientation.

Examining the free energy profiles of dimerization obtained for the N-terminal fragments of DISC1 with parallel (NN) and anti-parallel orientations (NN’) we found no significant difference free energy gap between both dimeric forms (Fig. 7 C). This could be rationalized by the absence of the central and C-terminal domains which provide most of the conformational heterogeneity observed for the full-length protein (Fig. 2). These results suggest that structural disorder by itself (i.e. without conformational heterogeneity) is enough to distinguish the parallel and anti-parallel dimer conformations.

Similarly, the free energy profiles of dimerization computed for the C-terminal fragments of DISC1 shows no significant different between dimers with parallel (CC) and anti-parallel orientations (CC’) (Fig. 7 D), which again suggest that both structural disorder and conformational heterogeneity are required to form stable DISC1 dimers.

## Conclusions

In this paper we present a detailed molecular level study of the behaviour of DISC1 protein which is believed to rely on disorder to act as a major hub for regulatory pathways connected with neural development. Specifically, we have shed light on the role of conformational heterogeneity and local structural disorder in the oligomerization pathway of the full-length DISC1 and its N and C-terminal fragments. By carrying out extensive sampling with coarse grained molecular dynamics simulations using the predictive energy landscape based protein force field AWSEM we have uncovered complex interplay between intra- and inter-domain contact frustration which shapes the conformational energy landscapes of DISC1 dimers.

We found that the interplay between intra- and inter-domain contacts drives the monomeric subunits of DISC1 into adopting an heterogeneous conformational ensemble with collapsed and exposed surfaces. We then go on to show that conformational heterogeneity and disorder translate into disparate binding pathways leading to formation of collapsed, transiently bound and exposed dimeric structures. We found that collapsed dimeric states have marginal oligomerization affinity whereas exposed states lead to formation of stable dimers. By analyzing the N and C-terminal fragments, we uncover that the structural disorder localized in N-terminal region is a significant driver of dimerization. The C-terminal region on the other hand is responsible for generating conformational heterogeneity and entropically favored exposed states, which have higher propensity for forming dimers and most likely serve as stepping stones for formation of higher level oligomers. Our simulation also predict the existence of collapsed dimeric states, which may act deep kinetic traps and serve as a powerful regulatory mechanism for controlling the oligomerization pathways and ultimately function of DISC1. We believe that the ability of DISC1 to populate an heterogeneous conformational space with both ordered and disordered regions and large exposed surfaces area can be a generic mechanism to regulate protein-protein interactions and potentially to initiate liquid-liquid phase separations.

## Acknowledgements

DAP acknowledges financial support from Caldwell Foundation of Iowa State.

