## Supporting Information for "Disorder mediated oligomerization of DISC1 proteins revealed by coarse-grained computer simulations"

### Supporting Information for: ”Disorder mediated oligomerization of DISC1 proteins studied by coarse-grained computer simulations.”

Authors: Julien Roche, Davit A. Potoyan

June 5, 2019

#### 1 Supporting Information

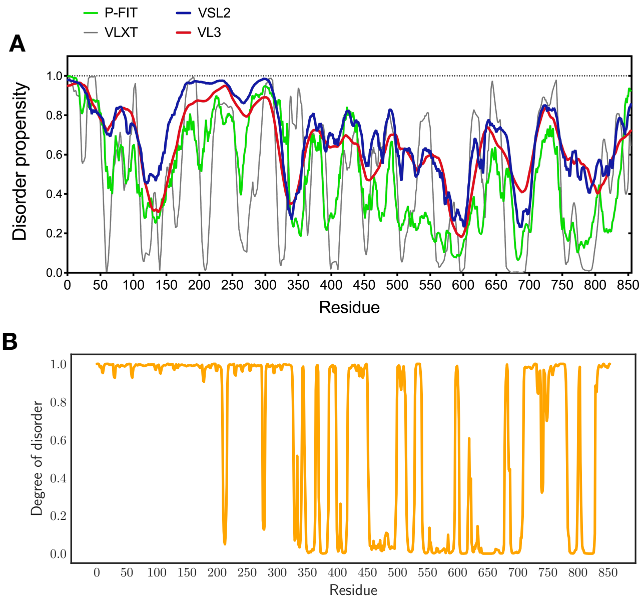


Figure 1: Quantifying disorder in DISC1 sequence. Shown are (A) A measure of disorder predicted based on ... (B) DSSP prediction converted to ’Degree of disorder” binary measure (C=0; H,E,I=1) obtained by averaging over ensemble of conformations generated in const T molecular dynamics simulations of full length DISC1 monomer.

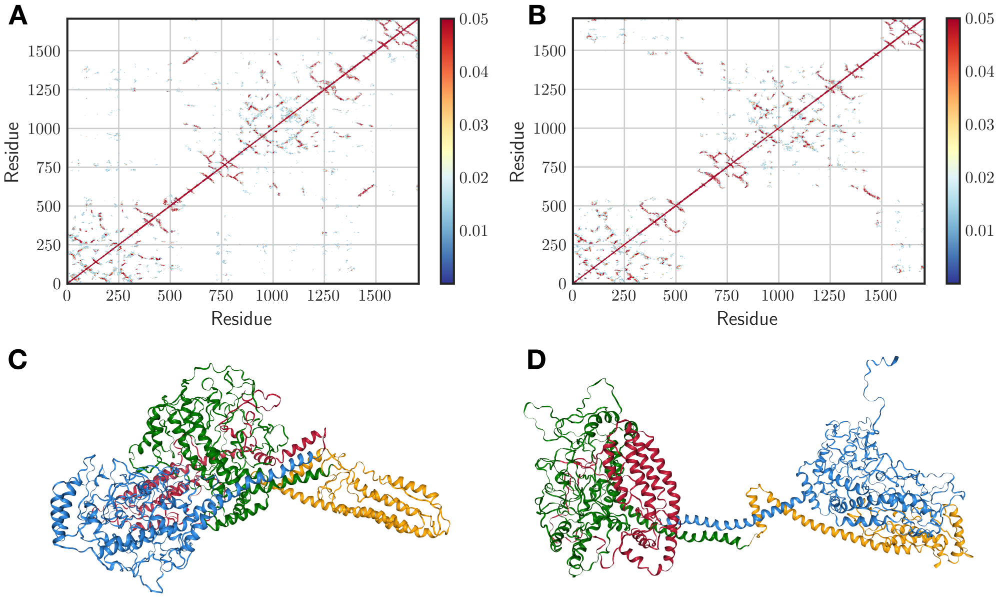


Figure 2: Shown are contact frequency map for full length DISC1 dimers (A,B) with representative conformations shown below (C,D)

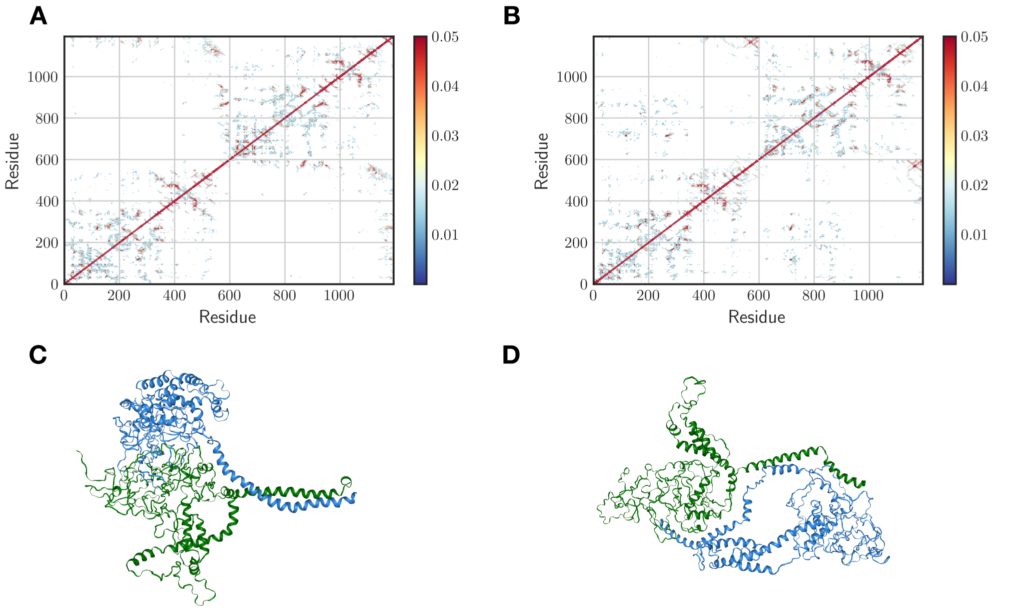


Figure 3: Shown are contact frequency map for N-terminal fragment DISC1 dimers (A,B) with representative conformations shown below (C,D)

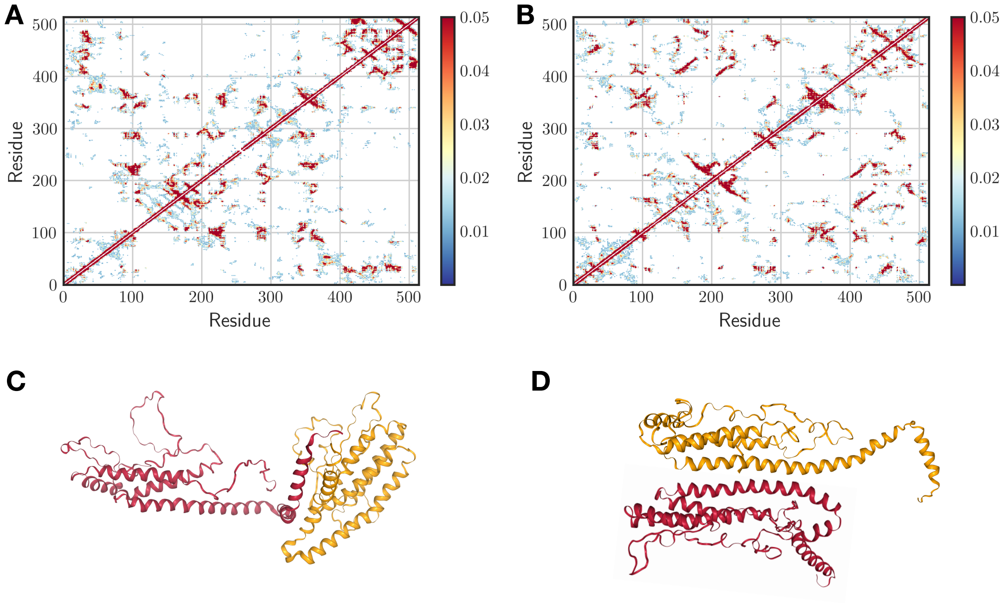


Figure 4: Shown are contact frequency map for C-terminal fragment DISC1 dimers (A,B) with representative conformations shown below (C,D)
